# Non-invasive approach for endoluminal biopsy coupled with single-cell proteomics allows for immune characterization of intracranial aneurysms

**DOI:** 10.1101/2025.02.17.638658

**Authors:** J. Antonios, B. Gultekin, B. Theriault, K. Yalcin, D. Miyagishima, N. Adenu-Mensah, N. Sujijantarat, A. Koo, J. Haynes, P. Cedeno, M. Johnson, R. Hebert, C. Matouk, T. Barak, M. Gunel

## Abstract

The immune regulatory mechanisms driving the stability, growth, and rupture of intracranial aneurysms (IAs) remain incompletely understood. In this study, we employ endoluminal biopsy with single-cell proteomics to comprehensively profile the immune landscape of IAs across their pathologic states. Our findings reveal distinct immune signatures associated with aneurysm states. Stable, i.e. non-growing unruptured, IAs exhibit a balanced immune cell composition. Ruptured IAs are marked by significant neutrophil predominance. Notably, we highlight key immune markers in aneurysm instability, offering new insights into immune drivers of aneurysm progression. These findings provide a foundation for immune-targeted, non-invasive therapeutic strategies aimed at targeting IAs and preventing rupture.

## Main

Saccular intracranial aneurysms (IAs) are localized dilatations of the cerebral arterial wall, affecting approximately 3% of the general population^1,2^. Annually, around 1.5% of IAs rupture ^2,3^, leading to subarachnoid hemorrhage (SAH), a severe complication characterized by high morbidity and mortality ^1,4-9^. The mechanisms driving IA formation and rupture remain incompletely understood, involving a complex interplay of environmental, hemodynamic, and genetic factors^2,10-14^. In the past decade and a half, substantial progress has been made in elucidating the genetic risk factors underlying IA through advanced high-throughput sequencing technologies ^12,15-21^. These efforts have identified multiple genetic loci and candidate genes associated with IA development ^22,23^. However, despite these advances, crucial aspects of IA biology remain underexplored, particularly the immune and tissue-level changes that dictate vessel wall stability or the progression toward rupture. Previous studies have established that aneurysm pathogenesis has an inflammatory component ^14,24-26^. Despite this, comprehensive immune profiling of IAs, particularly in relation to their radiographic stability or rupture risk, has been limited. In this study, we employ an innovative, non-invasive approach to map the immune landscape of IAs. Leveraging endoluminal biopsy techniques ^27-33^ and single-cell proteomics using mass cytometry by time-of-flight (CyTOF) ^34-37^, we present an extensive immune profiling of the largest high-fidelity IA patient cohort to date. Our findings reveal distinct immune signatures associated with aneurysm stability and rupture, providing critical insights that may pave the way for novel, immune-targeted, non-invasive strategies in early IA management.

To define the immune landscape of IAs, we optimized the technique for endoluminal IA biopsy, enabling the efficient collection of high-quality, high-yield samples from multiple patients. This approach allowed us to standardize the process, ensuring the integrity of the samples for subsequent analysis. Following this, we immune profiled specimens collected from 15 patients presenting with varied clinical scenarios during endovascular treatment. Study patients were identified prior to or at time of endovascular treatment, and samples were collected and preserved using cell preservation techniques for batch processing and analysis as described previously ^27-33^ (**Fig. 1a**). Clinical and radiographic data were recorded as outlined in the **Methods** (**Fig. 1b**, **Supp. Tables 1-3**). To characterize the immune environment of these IAs, we performed single-cell proteomic profiling using mass cytometry by time-of-flight (CyTOF). A 30-marker CyTOF panel (MaxPar, Standard Biotools, San Francisco, CA) was employed to identify major immune populations, including leukocytes (CD45), granulocytes (CD66b), B cells (CD19, CD20, IGD), T cells (CD3, CD4, CD8, TCRγδ), Natural Killer (NK) cells (CD16, CD56, CD57, CD161), and monocytes and subtypes (CD14, CD11c, HLA-DR). Additional markers for co-stimulation (CD27, CD28, IL7R [CD127]), activation (CD25, CD38, IL3R [CD123]), memory/naïve differentiation (CD45RO, CD45RA), and chemokines (CCR4, CD196 [CCR6], CCR7, CD183 [CXCR3], CXCR5), as well as the chemoattractant receptor CD294 [CRTH2], were also analyzed. Following stringent quality control (QC) steps as published previously^34,38^, 15 high-quality samples were retained for further analysis (**Methods**). Radiographic features were quantified, and using FlowJo (FlowJo, Ashland, OR) and custom Python analysis packages ^39-41^, we found that IA immune profiles segregated primarily into two populations: CD66b_High_CD45_Low_ and CD66b_Low_CD45_High_. Clustering of manually gated cells revealed 11 main populations, which were subsequently annotated as CD4+ T cells, CD8+ T cells, γδ T cells, B cells, dendritic cells, eosinophils, NK cells, mucosal-associated invariant T (MAIT)/ NK T cells, monocytes/macrophages, basophils, and neutrophils (**Fig. 2a-c**, **Supp. Fig. 2**).

**Figure 1.**
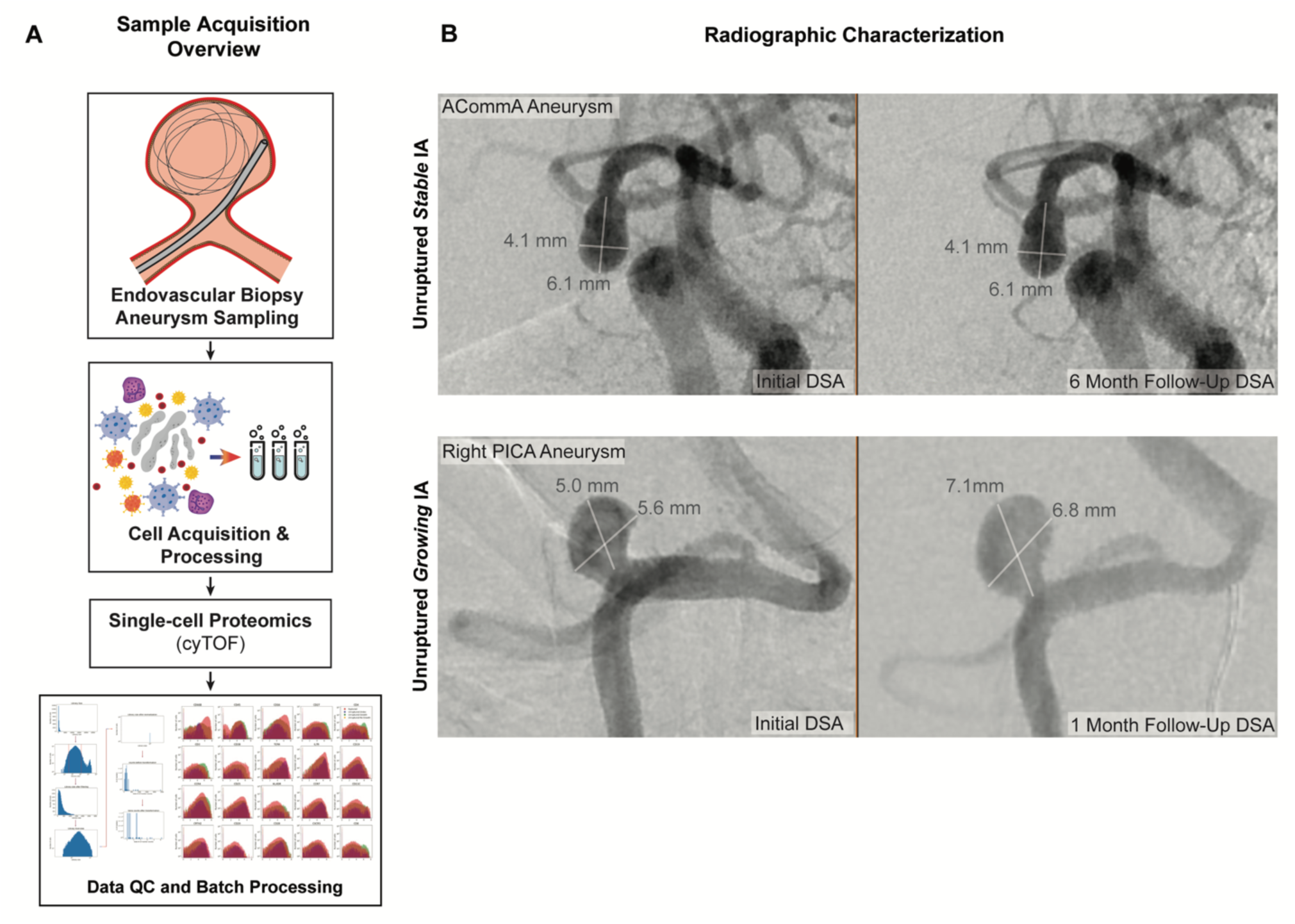
Sample acquisition and processing overview. **(A)** Endoluminal biopsy of intracranial aneurysm (IA) was performed via cell acquisition from discarded coils deployed and then removed from aneurysm, as well as microcatheter tips. Cells are processed along with red blood cell (RBC) lysis and dissociation. This was followed by quality control and sub-processing for single cell mass cytometry time-of-flight (cyTOF). **(B)** An example of an unruptured stable IA (anterior communicating artery (ACommA) aneurysm) is shown with stable measurements of 4.1 x 6.1 mm at initial and follow-up digital subtraction angiography (DSA) studies. This is followed by an example of an unruptured growing IA (right posterior inferior cerebellar artery (PICA) aneurysm) with growth documented at initial (5.0 x 5.6 mm) and follow-up (7.1 x 6.8 mm) DSAs.

**Figure 2.**
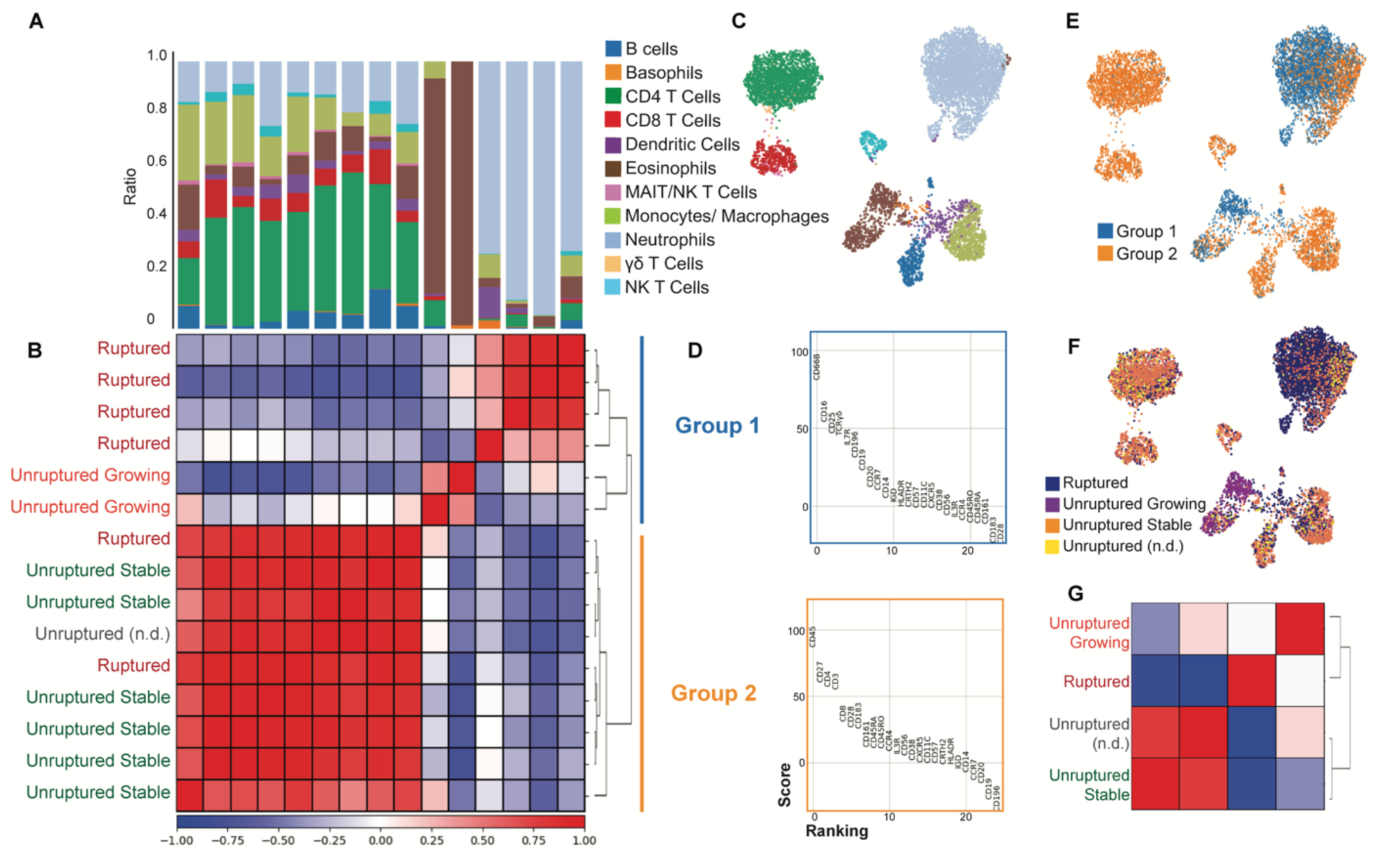
Single-cell proteomics demonstrates distinct immune clusters. Cell populations were identified based on their top expressed markers as follows: monocytes, immature granulocytes, neutrophils, CD8 T cells, CD4 T cells, NK cells, and B cells. **(A)** Immune cell ratios are demonstrated on this plot, showing shifts in cells across patient subsets. **(B)** Pearson correlation plot demonstrates two distinct groups of IAs, annotated with labels for aneurysm state and sub-clusters (Group 1 and Group 2). **(C)** The representation of immune cells is demonstrated on the UMAP plot. **(D)** The differential expression plots showing the highest upregulated markers demonstrate distinct differences between Group 1 and Group 2 clusters. **(E)** The representation of cyTOF immune clusters Group 1 and Group 2 is demonstrated on the UMAP plot. **(F)** The representation of IA state (ruptured, unruptured growing, unruptured stable, and unruptured (not documented) (n.d.)) is demonstrated on the UMAP plot. **(G)** Correlation plot shows that the immune profiles of unruptured growing IAs and ruptured IAs are statistically similar while unruptured stable IAs cluster separately.

Hierarchical clustering grouped samples into two distinct clusters based on their immune profiles (**Fig. 2a-d**). The first, Group 1, consisted only of unstable aneurysms, defined as ruptured IAs (n = 4) and unruptured growing IAs (n=2) (aneurysms with a documented diameter increase on serial imaging using the same modality). Group 2 was comprised predominantly of stable aneurysms, defined as unruptured stable IAs (n = 6) (no growth with stable diameter on serial imaging using the same modality). Group 2 also included two ruptured IAs and one with undocumented stability. We assessed clinical and radiographic variables to identify potential drivers of these immune signatures. We found that there was a significant association between cyTOF immune groups and IA stability (Fisher Exact test, two-tailed, p=0.0097, OR = Inf, 95% CI 1.968 to Inf) (**Supp. Table 4**). Interestingly, no other significant associations were observed between the immune groups and other variables, such as IA laterality (p=0.6224), hypertension (p=0.6224), or smoking status (p=0.6224) (**Supp. Table 4**; **Supp. Fig. 3a-d**). Of note, distinct cell populations were enriched in Group 1 when compared to Group 2 (**Fig. 2e**), which was also recapitulated when examining cell populations across IA stability states (**Fig. 2f**). This suggested that there was a correlation between specific cell populations and corresponding IA stability states. We found that the immune profiles of unruptured growing and ruptured IAs clustered together, while unruptured stable IAs formed a separate cluster (**Fig. 2g**). Interestingly, ruptured IAs exhibited significant neutrophil enrichment. The immune cells in unruptured IAs were more heterogenous. In summary, the data indicate that CyTOF immune group stratification may offer insight into IA stability, whereas other examined clinical and demographic factors appear to be less relevant in this context.

Given the enrichment of neutrophils in ruptured IAs, we next asked whether the expression profiles of neutrophils differed across IA rupture states. We found that activation marker CD38, which has been shown to regulate neutrophil chemotaxis^42-49^, was upregulated on neutrophils in ruptured IAs. We also identified a significant increase in expression of HLA-DR, a surface molecule that neutrophils express in response to inflammatory cytokines to acquire accessory cell functions that help activate T cells ^50,51^, in this population (**Fig. 3a, b**; **Supp Fig. 4a-d**). Additionally, the chemokine receptor CCR7, associated with neutrophil migration to lymph nodes and subsequent activation^52^, was enriched in neutrophils in ruptured IAs (**Fig. 3c**). Unruptured growing IA neutrophils highly expressed CD28, which regulates migration by modulating CXCR1 expression^53^, in addition to CXCR3 and CXCR5 ^54-57^. Unruptured stable IA neutrophils expressed CCR6, a marker that has been generally associated with angiogenesis ^58-62^. These findings collectively support a correlation between distinct neutrophil profiles and the rupture status of IA.

**Figure 3.**
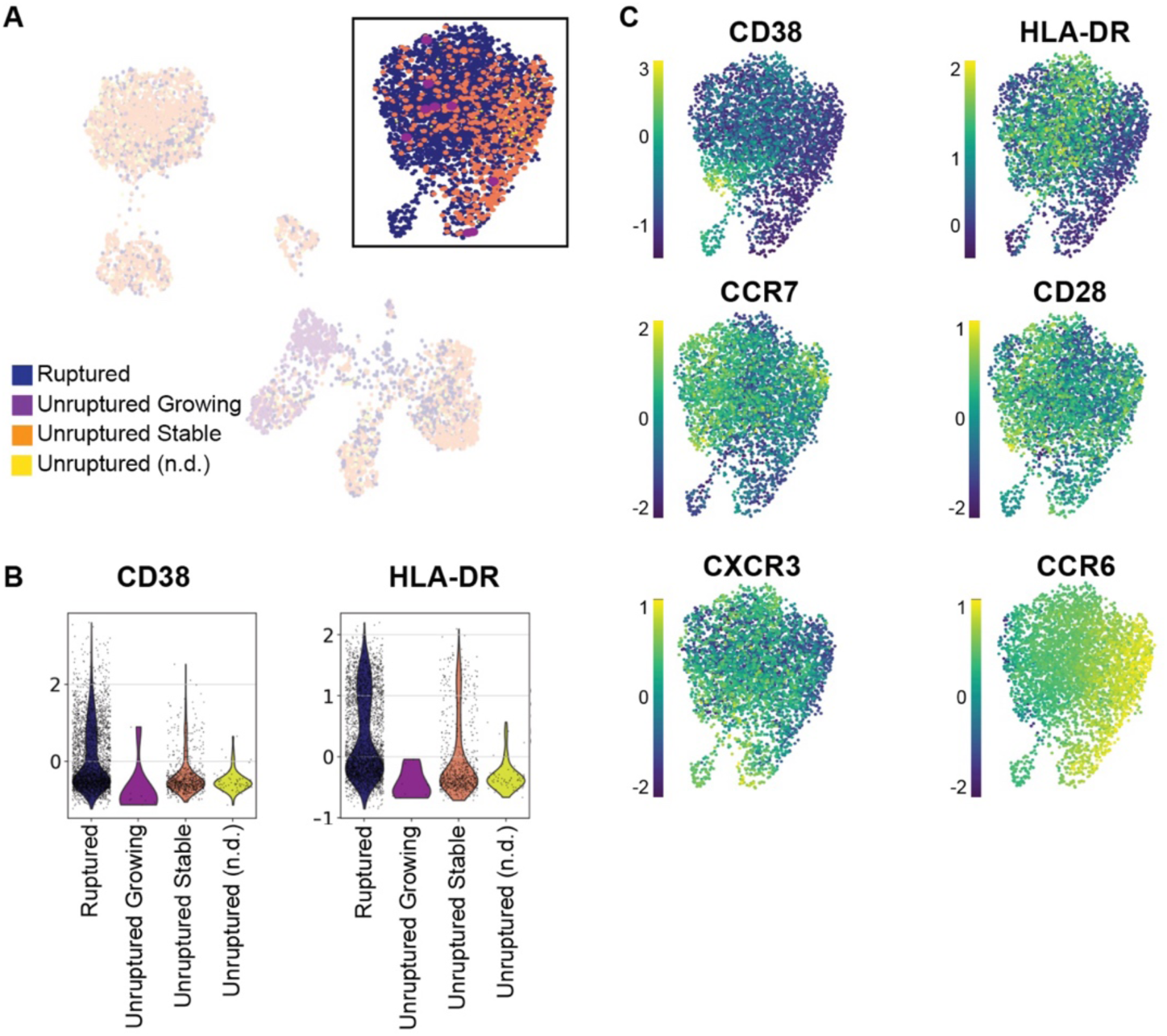
Neutrophil immune population shifts across IA rupture status. The representation of IA rupture status (ruptured, unruptured growing, unruptured stable, and unruptured (not documented) (n.d.)) is again demonstrated on the **(A)** UMAP plot with emphasis on the neutrophil population, with **(B)** quantification of CD38 and HLD-DR expression across them. **(C)** Expressions of relevant markers CD38, HLA-DR, CCR7, CD28, CXCR3, and CCR6 within the neutrophil population are visualized on UMAP plots

Overall, this is the first study to utilize endovascular sampling — a novel method for endoluminal biopsy — to comprehensively profile immune cells in human IA samples. Compared to prior human IA studies, which were limited by small sample sizes or a narrow focus on single patients, our approach represents a significant advancement. By employing single-cell mass cytometry (CyTOF), we generated the most extensive dataset of immune proteomic profiles in human IAs to date. Our study provides novel insights into the immune landscape of IAs across different states—stable, unruptured growing, and ruptured. We found that stable IAs exhibited a balanced immune cell composition and ruptured IAs were characterized by a significant neutrophil predominance. Although unruptured growing IAs showed an enrichment of eosinophils, the sample set was small and did not allow for further study. The decreased lymphocytic cell ratios observed in both unruptured growing and ruptured IAs further emphasize the shift from adaptive to innate immune responses as aneurysms progress. These distinct immune signatures suggest that immune landscape dynamics play a critical role in IA stability, growth, and rupture.

Our findings align with previous studies, which point to the critical role of neutrophils in aneurysm progression ^3,63-67^. The enrichment of neutrophils in ruptured IAs suggests that these cells play a pivotal role in aneurysm rupture state. Our findings support a model in which a baseline immune composition in stable IAs—comprising T cells, B cells, and myeloid cells—shifts in aneurysms that are growing. The marked neutrophil predominance in ruptured IAs could likely be secondary to dynamic changes in the immune mileu of the aneurysm preceding rupture or an acute post-hemorrhagic inflammatory response. This is consistent with previous studies and underscores the interplay between immune mechanisms and aneurysm progression^24^.

We acknowledge the limitations of our cross-sectional design, which restricts our ability to establish causality. Moreover, the logistical and ethical challenges of conducting longitudinal studies in IA patients pose an obstacle to further elucidating the temporal dynamics of immune infiltration. Despite these limitations, our findings open new avenues for developing targeted, non-invasive therapies that leverage immune modulation to stabilize aneurysms and prevent rupture. Future research utilizing advanced single-cell technologies could provide deeper insights into the cellular heterogeneity within IAs, offering the potential for personalized immune-based management strategies. For instance, identifying patient-specific immune profiles could help stratify aneurysm rupture risk and inform treatment decisions, paving the way for more precise interventions. In conclusion, our study provides compelling evidence of the immune system’s central role in the development and rupture of intracranial aneurysms.

## Online Methods

### Patient and Aneurysm Characterization

We collected information regarding co-morbidities (smoking history, hypertension), family and personal history, clinical presentation (headache, radiographic findings including modified Fisher score, presence of a cranial nerve palsy), time to treatment from symptom onset and from rupture onset (where applicable), aneurysm location and laterality, aneurysm morphology (features, height, width, diameter, circumference in three dimensions, neck diameter, volume, aspect ratio, bottleneck factor, height-width ratio), and aneurysm growth rate where applicable ^68-70^. Aneurysm growth rate was specifically determined by measuring aneurysm diameter from two timepoints (the day of treatment and the most recently available imaging study prior) in the same plane and projection on digital subtraction angiography (DSA) or computerized topography angiography (CTA), and then calculating the change in size as a unit of time. The radiographic measurements were validated independently by two neuroradiologists and one neurosurgeon. These parameters allowed us to fully quantify all of the factors associated with our data set in the subsequent analyses.

### Endoluminal Biopsy

Endoluminal biopsy was performed as previously described ^27-29,32,33,71^, with sample acquired at time of treatment. Our approach utilized two primary sources of cellular material: firstly, coils that were initially deployed within the aneurysm but not permanently attached, allowing for their subsequent retraction; and secondly, microcatheter tips, which were positioned inside the aneurysm throughout the treatment duration and removed. In both cases, cells were washed in PBS x1 buffer followed by RBC Lysis Buffer (eBioscience, ThermoFisher, San Diego, CA) and stored in -80C for batch processing using cell freezing media (ThermoFisher, San Diego, CA).

### Mass cytometry

In this study, we implemented the CyTOF workflow using the Maxpar Direct Immune Profiling Assay (Standard Biotools, San Francisco, CA), which is designed to identify a spectrum of immune cell populations with a 30-marker antibody panel. The antibodies used in the study were labeled with various isotopes and include CD45 (HI30)-89Y, CD196/CCR6 (G034E3)-141Pr, CD123 (6H6)-143Nd, CD19 (HIB19)-144Nd, CD4 (RPA-T4)-145Nd, CD8a (RPA-T8)-146Nd, CD11c (Bu15)-147Sm, CD16 (3G8)-148Nd, CD45RO (UCHL1)-149Sm, CD45RA (HI100)-150Nd, CD161 (HP-3G10)-151Eu, CD194/CCR4 (L291H4)-152Sm, CD25 (BC96)-153Eu, CD27 (O323)-154Sm, CD57 (HCD57)-155Gd, CD183/CXCR3 (G025H7)-156Gd, CD185/CXCR5 (J252D4)-158Gd, CD28 (CD28.2)-160Gd, CD38 (HB-7)-161Dy, CD56/NCAM (NCAM16.2)-163Dy, TCRγδ (B1)-164Dy, CD294 (BM16)-166Er, CD197/CCR7 (G043H7)-167Er, CD14 (63D3)-168Er, CD3 (UCHT1)-170Er, CD20 (2H7)-171Yb, CD66b (G10F5)-172Yb, HLA-DR (LN3)-173Yb, IgD (IA6-2)-174Yb, CD127 (A019D5)-176Yb, and Live/dead intercalator-103Rh-103Rh. These metal-conjugated antibodies were employed in the study to perform high-parameter single-cell mass cytometry, enabling the simultaneous analysis of various cellular markers at the single-cell level ^39,72,73^. Cells were labeled and fixed ^41,74^. Afterwards, samples were assessed by the CyTOF2 (Fluidigm) using a flow rate of 0.045 mL/min in the presence of EQ Calibration beads (Fluidigm) for normalization as previously described^34^.

Samples underwent gating steps as described using the MaxPar (Standard Biotools, San Francisco, CA) gating protocol ^75-78^. After discarding debris as well as data that did not gate into morphologic and cellular gates, the remaining high-quality samples were further selected for by excluding samples with <100 cells for a total sample size of 15. This was performed in order to analyze only high integrity samples with higher counts of cells. Analysis was performed as follows: initial gating for data cleanup was performed on the FlowJo platform using the MaxPar gating strategy by exclusion of debris (Iridium^low^, DNA^low^), multicell events (Iridium^hi^, DNA^hi^), and dead cells (cisplatin^hi^). CyTOF data was then imported into a Python-based tool and then analyzed using Principal Component Analysis, uniform manifold approximation and projection (UMAP), followed by Leiden clustering to delineate cell populations.

### Data and Statistical Analysis

We analyzed both our cyTOF data using techniques extensively described and documented previously ^34,38,79-81^. Statistical analysis was performed using Prism (version 10.9.0). Pairwise Fisher exact tests were performed to assess significance with respect to patient, clinical, and radiographic variables against immune cell populations. The significance level in all tests was P < 0.05.

## Reporting summary

Further information on research design is available in the Nature Portfolio Reporting Summary linked to this article.

## Data availability

All source data are provided in the supplementary materials; additional details are available from the authors upon reasonable request.

## Code availability

The code used in this study is available from GitHub (link).

## Figures

**Supplementary Figure 1.**
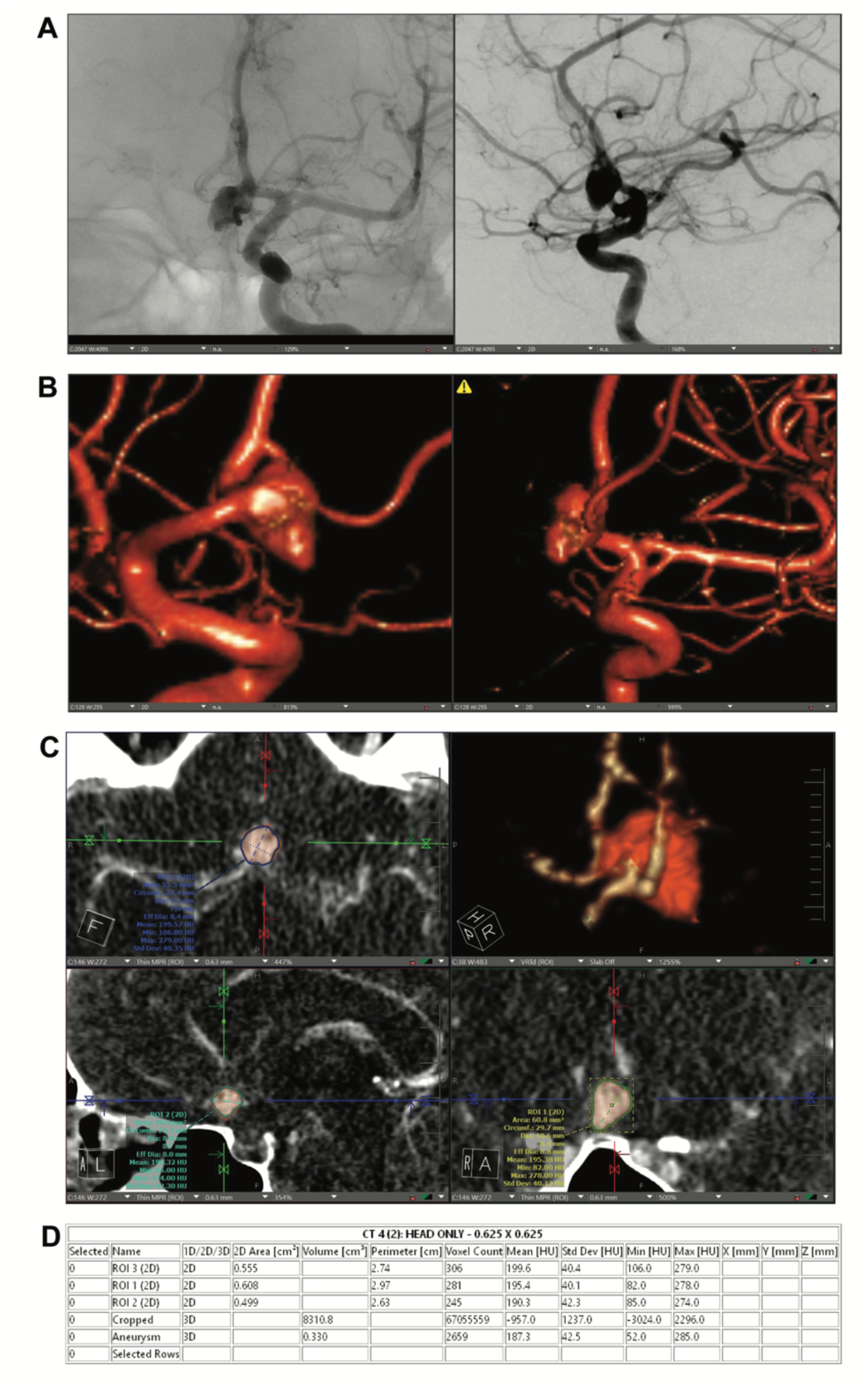
Example pipeline for quantitation of aneurysm radiographic parameters. **(A)** Digital subtraction angiography demonstrates the aneurysm in A/P and lateral planes with **(B)** 3D reconstruction delineating the morphologic features of the aneurysm. **(C)** This is recapitulated on computed topography angiography (CTA) imaging and 3D reconstruction with measurements and **(D)** tabulations of individual radiologic parameters.

**Supplementary Figure 2.**
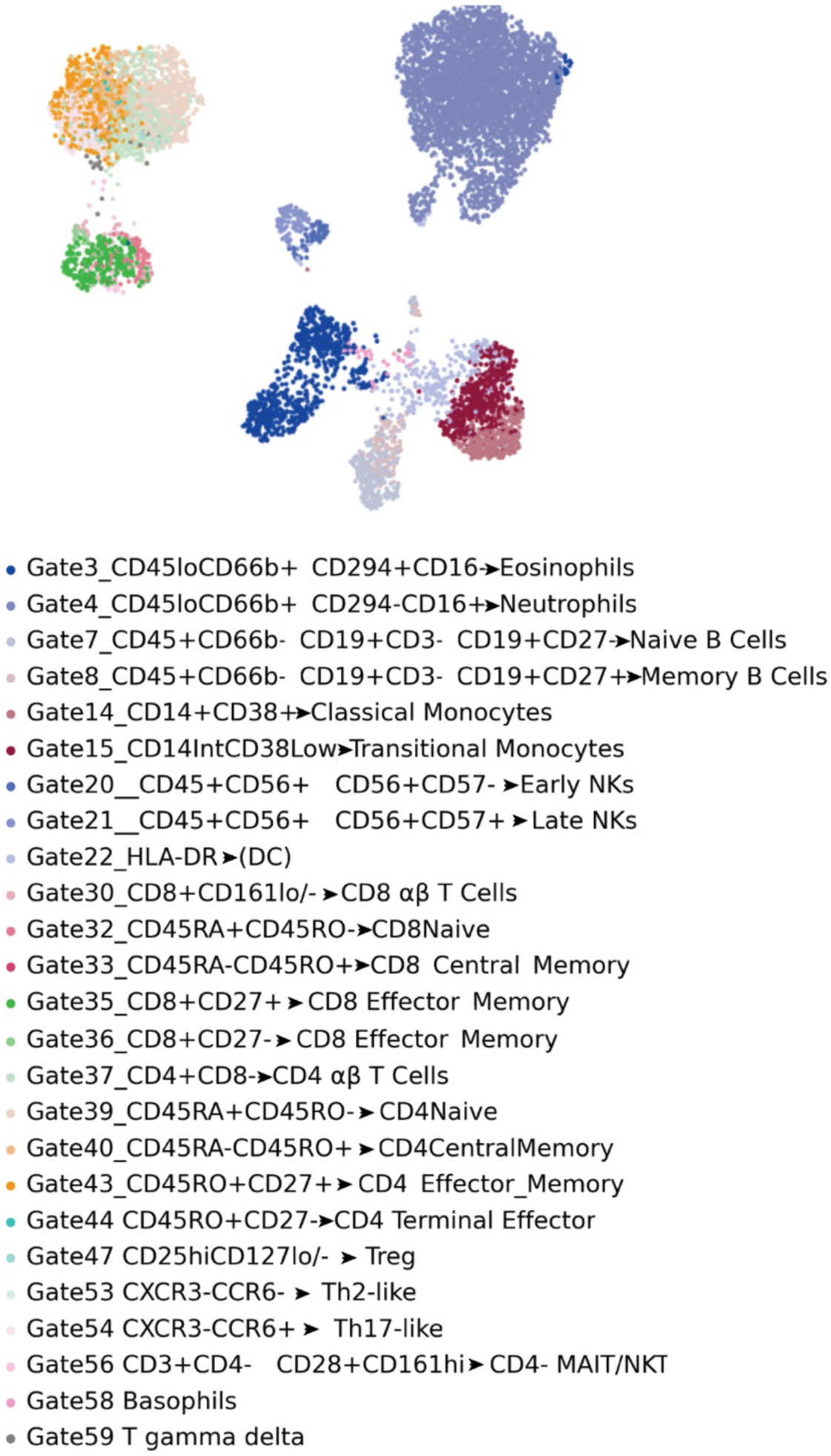
Manual gating corroborates unbiased clustering. We performed manual gating using the cell-labeling rubric provided with the cyTOF kit ^78^. The cell populations identified corroborated the unbiased clustering and labeling performed.

**Supplementary Figure 3.**
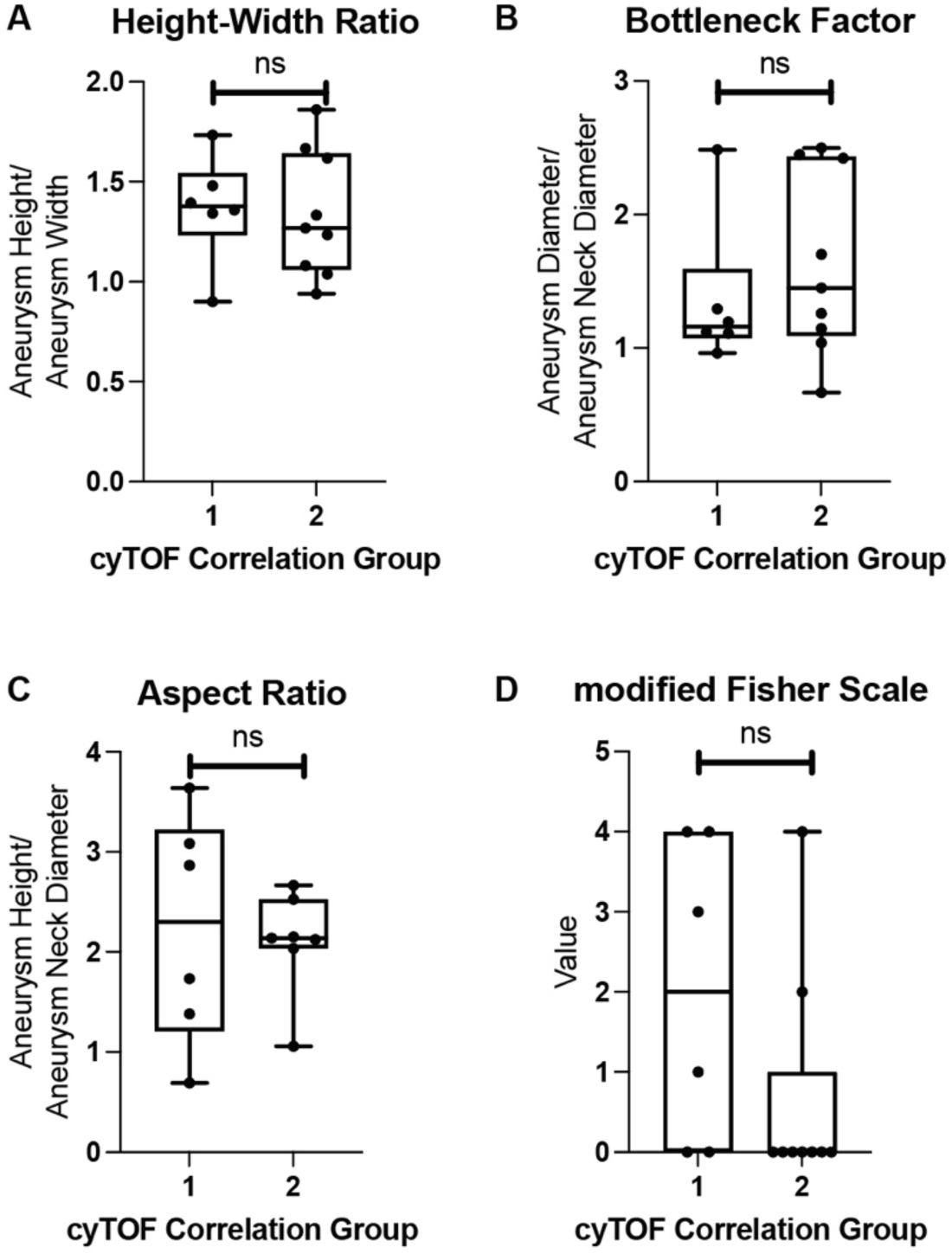
Relationship between clinical/radiological features and the cyTOF findings. The representation of **(A-C)** morphologic (aspect ratio, bottleneck factor, and height-width ratio) and **(D)** patient parameters (modified Fisher Scale, across each cyTOF immune cluster (Group 1 and Group 2) were plotted, showing no clear contribution to any immune signature from any parameter.

**Supplementary Figure 4.**
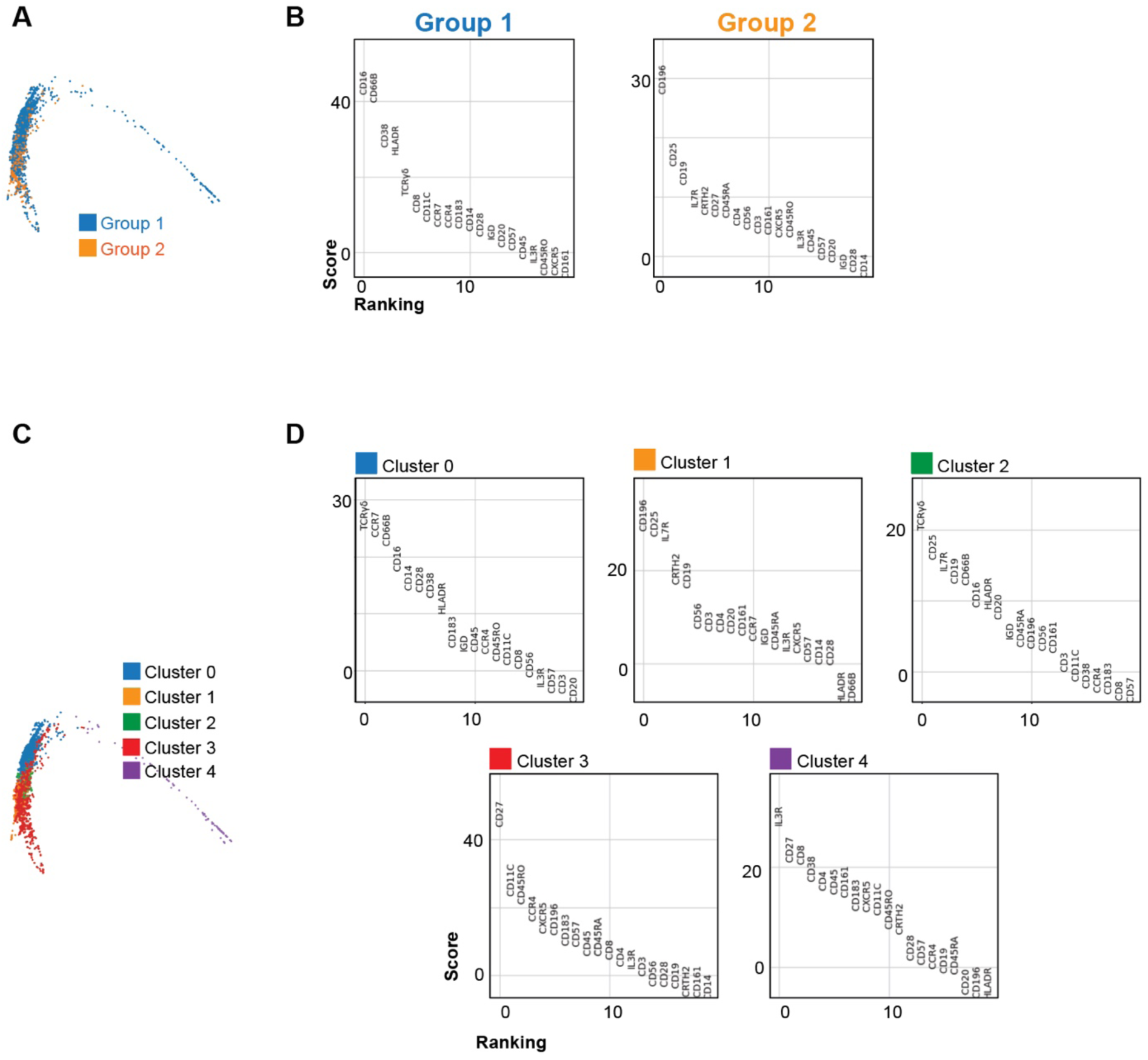
Differential expression changes across neutrophils. **(A)** Differential expression analysis was performed with PHATE plot visualization showing shifts in the neutrophil population across the Group 1 and Group 2 sub-clusters. **(B)** Differential expression plots for neutrophils in Group 1 and Group 2 show the markers with the highest expression. **(C)** We performed Leiden sub-clustering of the neutrophil population and noted that there were 5 distinct sub-clusters of neutrophils (Cluster 0 – 4) visualized with PHATE plot showing shifts across the clusters. **(D)** Differential expression plots showing the markers with the highest expression are shown below.

## Supplementary Tables

**Supplementary Data Table 1.** Overview Patient Characteristics. The age, sex, smoking history, and hypertension history of the patients is represented here.

| **Variable** |  | *Total (n=15)* |
| --- | --- | --- |
| *Age* |  |  |
|  | Mean (SD) | 52.20 (12.42) |
|  | Range | 28 – 67 |
| *Sex* |  | *Count (%)* |
|  | Female | 8 (53.33%) |
|  | Male | 7 (46.67%) |
| *Smoking History* |  |  |
|  | Current | 4 (26.67%) |
|  | Former | 5 (33.33%) |
|  | Never | 6 (40.00%) |
| *Hypertension History* |  |  |
|  | Controlled | 4 (26.67%) |
|  | Poorly Controlled | 4 (26.67%) |
|  | No Hypertension | 7 (46.67%) |
| *Sample Viability* |  |  |
|  | Mean (SD) | 93.28 (5.11) |
|  | Range | 78.83 – 98.08 |

**Supplementary Data Table 2.**
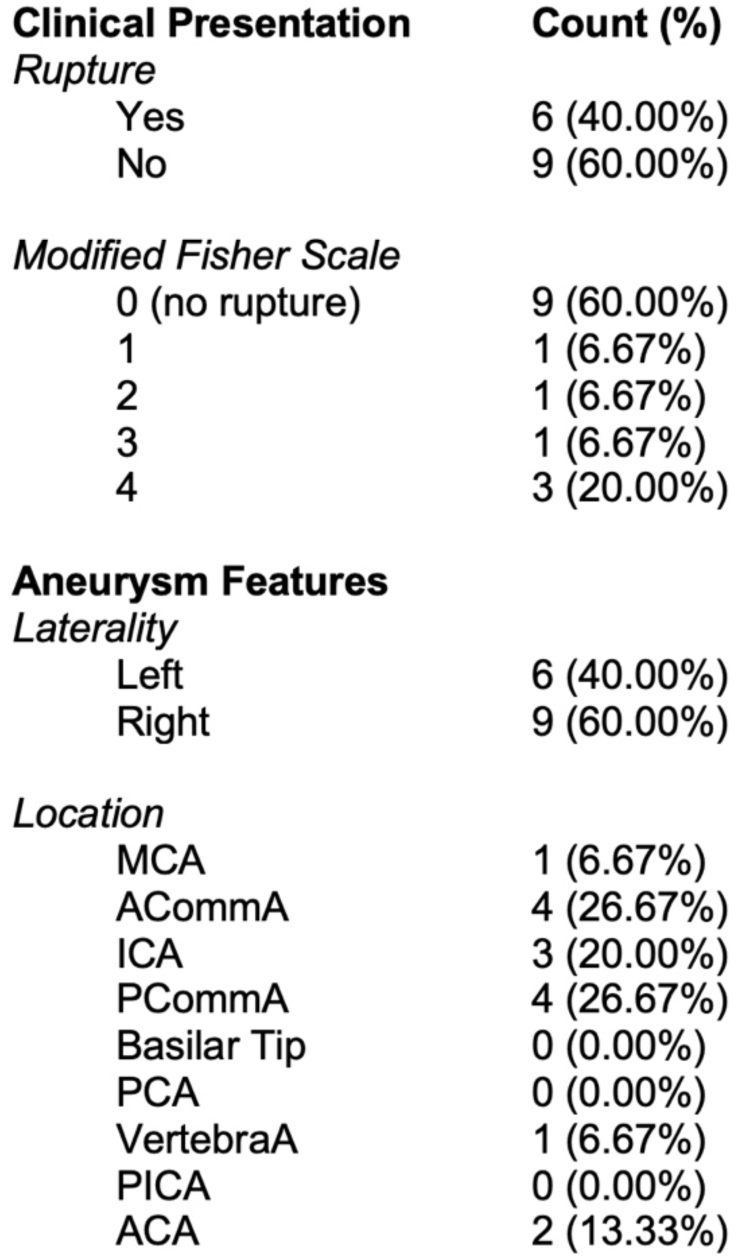
Overview of Patient Presentation and Aneurysm Location. The clinical presentation, including rupture status, modified Fisher Scale on computed topography (CT) imaging, as well as aneurysm laterality and location, are shown here.

**Supplementary Data Table 3.** Aneurysm Morphology Characterization. The aneurysm morphologies documented are summarized here, including aneurysm height, width, diameter, neck diameter, volume, aspect ratio, bottleneck factor, and height-width ratio.

|  |  |
| --- | --- |
| **Measurement** |  |
| *Aneurysm Height (H)* | (mm) |
| Mean (SD) | 7.4 (4.2) |
| Range | 1.0 – 22.8 |
| *Aneurysm Width (W)* |  |
| Mean (SD) | 6.0 (4.6) |
| Range | 0.9 – 27.2 |
| *Aneurysm Diameter (D)* |  |
| Mean (SD) | 6.8 (4.9) |
| Range | 1.1 – 23.3 |
| *Neck Diameter (nD)* |  |
| Mean (SD) | 3.8 (1.2) |
| Range | 1.9 – 6.4 |
| *\*Volume (V)* | (mm<sup>3</sup>) |
| Mean (SD) | 2.3 (3.9) |
| Range | 0.3 – 15.5 |
| *Aspect Ratio (H/nD)* |  |
| Mean (SD) | 2.1 (0.7) |
| Range | 0.7 – 3.6 |
| *Bottleneck Factor (H/D)* |  |
| Mean (SD) | 1.5 (0.6) |
| Range | 0.5 – 2.8 |
| *Height-Width Ratio (H/W)* |  |
| Mean (SD) | 1.3 (0.4) |
| Range | 0.8 – 2.4 |

**Supplementary Data Table 4.** Fisher Exact tests and odds ratios for pertinent categories. Table summarizing the results of Fisher Exact tests and odds ratios for associations between categorical variables. Each row corresponds to a pair of variables tested, with columns showing p-value of Fisher Exact testing, as well as the odds ratio and the 95% confidence interval. Significant associations (p < 0.05) are noted in bold, highlighting meaningful relationships between variable pairs. (*One of our unruptured aneurysms did not have stability documented and was not included for this pairwise test.)

|  | cyTOF Group 1 | cyTOF Group 2 | p-value | OR | 95% CI |
| --- | --- | --- | --- | --- | --- |
| **# Patients** | 6 | 9 |  |  |  |
| **Stability Status (Unstable)\*** | 6 | 2 | 0.0097 | Inf | 1.968 to inf |
| **Rupture Status (Ruptured)** | 4 | 2 | 0.1357 | 7 | 0.6280 to 51.85 |
| **Laterality (Left)** | 3 | 3 | 0.6224 | 0.5 | 0.07776 to 3.378 |
| **HTN (Present)** | 3 | 6 | 0.6224 | 2 | 0.2960 to 12.86 |
| **Smoking (Current/Former)** | 3 | 6 | 0.6224 | 0.5 | 0.07776 to 3.378 |

